# Impact of DNA sequencing and analysis methods on 16S rRNA gene bacterial community analysis in dairy products

**DOI:** 10.1101/305078

**Authors:** Zhengyao Xue, Mary E Kable, Maria L Marco

## Abstract

DNA sequencing and analysis methods were compared for 16S rRNA V4 PCR amplicon and gDNA mock communities encompassing nine bacterial species commonly found in milk and dairy products. The communities were examined using Illumina MiSeq and Ion Torrent PGM DNA sequencing methods followed by the QIIME 1 (UCLUST) and Divisive Amplicon Denoising Algorithm 2 (DADA2) data analysis pipelines including taxonomic comparisons to the Greengenes and Ribosomal Database Project (RDP) databases. Examination of the PCR amplicon mock community with these methods resulted in Operation Taxonomy Units (OTUs) and Amplicon Sequence Variants (ASVs) that ranged from a low of 13 to high of 118 and were dependent on the DNA sequencing method and read assembly step. The elevated numbers of OTUs and ASVs included assignments to spurious taxa as well as sequence variants of the nine species included in the mock community. Comparisons between the gDNA and PCR amplicon mock communities showed that combining gDNA from the different strains prior to PCR resulted in up to 8.9-fold greater numbers of spurious OTUs and ASVs. However, the DNA sequencing method and initial data assembly steps conferred the largest effects on predictions of bacterial diversity, independent of the mock community type (PCR amplicon or gDNA; Bray-Curtis R^2^ = 0.88 and weighted Unifrac, R^2^ = 0.32). Overall, DNA sequencing performed with the Ion Torrent PGM and analyzed with DADA2 and the Greengenes database resulted in the most accurate predictions of the mock community phylogeny, taxonomy, and diversity.

**Importance:** Validated methods are urgently needed to improve DNA-sequence based assessments of complex bacterial communities. In this study, we used 16S rRNA PCR amplicon and gDNA mock community standards, consisting of nine, dairy-associated bacterial species, to evaluate the most commonly applied 16S rRNA marker gene DNA sequencing and analysis platforms used in evaluating dairy and other bacterial habitats. Our results show that bacterial metataxonomic assessments are largely dependent on the DNA sequencing platform and read curation method used. DADA2 improved sequence annotation compared with QIIME 1, and when combined with the Ion Torrent PGM DNA sequencing platform and the Greengenes database for taxonomic assignment, the most accurate representation of the dairy mock community standards was reached. This approach will be useful for validating sample collection and DNA extraction methods and ultimately investigating bacterial population dynamics in milk and dairy-associated environments.

## Introduction

Advancements in massively-parallel, DNA sequencing technologies have resulted in a dramatic increase in knowledge of the microorganisms found in natural environments, food systems, and the human body. 16S rRNA gene amplicon sequencing, in particular, has been a cornerstone for investigating bacterial diversity and phylogeny. This approach has enabled the simultaneous identification of the majority of bacteria in complex microbial communities. Although analysis of 16S rRNA gene diversity has provided significant new perspectives on bacterial habitats, there remain challenges to sample preparation, DNA sequencing, and data analysis approaches for ensuring accurate measurements of bacterial populations.

To address these issues, recent studies have compared sample collection methods (1–3) and storage conditions (2, 4–9). These studies generally showed that differences in bacterial composition caused by those methodological alterations are relatively minor compared to intersample variation (1, 3, 5–9). DNA extraction methods, on the other hand, can result in major changes to estimates of bacterial proportions between Gram-positive and Gram-negative bacteria which are more or less difficult to lyse (4, 7, 10–15). Moreover, PCR can also introduce bias depending on the DNA polymerase (16), number of cycles (17) and variable region of the 16S rRNA gene being compared (4, 18, 19).

DNA sequencing platforms, including 454 pyrosequencing, Illumina, Ion Torrent and Pacific Biosciences have also been shown to cause variation in the final community assessments (4, 18–21). However, some of studies also employed different post-run strategies (4, 20), rendering the relative influence of DNA sequencing and read assembly and trimming unclear on the outcomes. Moreover, data analysis methods, especially read clustering approaches (i.e. generating representative sequences), are known to have a significant impact on the interpretation of bacterial composition (21–27). Reference-based sequence clustering tends to underestimate community diversity because it is limited by the sequences included in the reference database (21, 23, 27). *De novo* sequence clustering can recover more sequence variants than reference-based methods, but can also result in unstable Operational Taxonomic Units (OTUs) between projects that are composed of different sequences with each clustering iteration at the 97% nucleotide sequence identity threshold (28, 29). As a result, there is now effort to move away from OTU based methods towards DNA sequences that represent single nucleotide variation (30, 31). One of the amplicon sequence variants (ASVs) clustering methods is DADA2 (Divisive Amplicon Denoising Algorithm 2), which builds a quality-based model for filtering error and identifying variation in 16S rRNA gene sequences (26).

Herein, we sought to compare different DNA sequencing and data analysis strategies for the capacity to accurately detect the composition of a mock bacterial community consisting of nine species commonly found in milk. To eliminate biases introduced by sample type and DNA extraction method, we employed two different mock communities consisting of either organism-specific PCR amplicons or purified genomic DNA (gDNA). This approach allowed us to compare the performance of two popular benchtop DNA sequencers (Illumina MiSeq and Ion Torrent PGM), sequence read assembly, and OTU/ASV analysis methods.

## Results

### Comparison of different DNA sequence analysis methods for 16S rRNA V4 region reads generated with the Illumina MiSeq and Ion Torrent PGM

A mock community was prepared by combining 16S rRNA V4 region PCR amplicons from nine bacterial strains in equimolar quantities prior to DNA sequencing on Ion Torrent PGM and Illumina MiSeq instruments. Sequences were either assembled (Illumina MiSeq) or maintained as single-end (Illumina MiSeq and Ion Torrent PGM) reads. A low percentage of reads were identified as chimeras (0 to 1.4 %) (Table S1) and the remaining reads were analyzed using QIIME 1 (UCLUST) or DADA2 pipelines for OTU or ASV identification using the Greengenes (version 13.8) and RDP (version GOLD for QIIME 1 and version 11.5 for DADA2) reference databases.

### Illumina MiSeq paired-end assemblies

QIIME 1 analysis of Illumina MiSeq paired-end, assembled reads with recommended parameters (32) resulted in total OTU numbers that were at least 4.2-fold greater than the expected nine OTUs encompassing strains included in the mock community (Table 2). When the Greengenes database was used for OTU alignment, 85 OTUs were identified. The majority of those OTUs (65) were assigned to taxa included in the mock community. Although the numbers of OTUs varied for each taxon, a single OTU representative encompassed the majority (> 60%) of reads for each of the mock community members (Table 2). For example, out of nine *Staphylococcus* OTUs identified with Greengenes, 99% of the reads were represented by one OTU. The remaining 20 OTUs identified with Greengenes were either designated as taxa that were not included in the mock community or were designated only to the order level. When the RDP database was used as the reference database for QIIME 1 analysis, the total OTUs decreased to 65, largely due to the reduction in spurious “other” assignments (Table 2). However, the overall OTU number remained considerably higher than the expected nine OTUs based on the mock community composition.

**Table 1.**
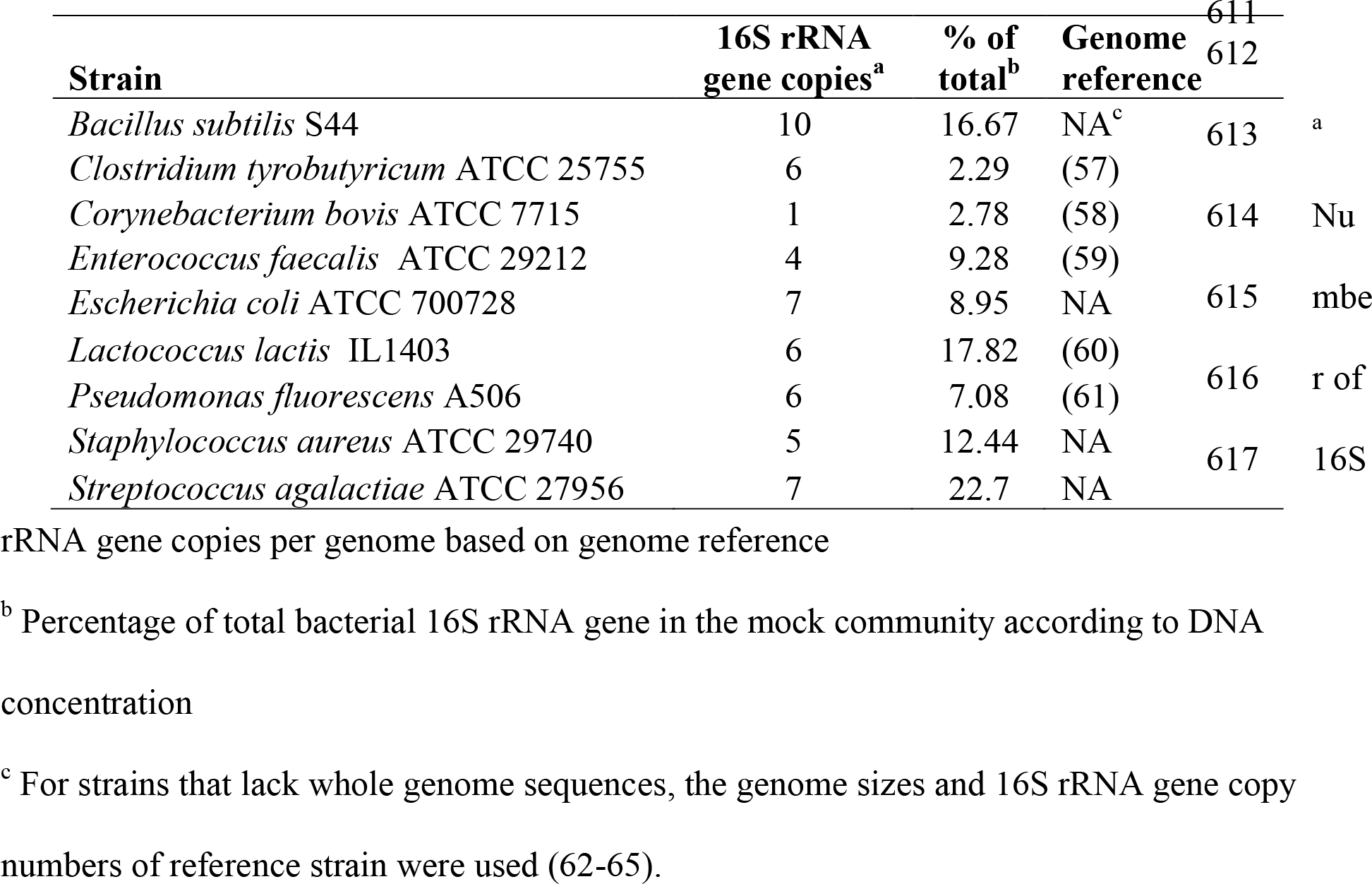
Bacterial strains and expected relative abundances in the gDNA mock community.

**Table 2.**
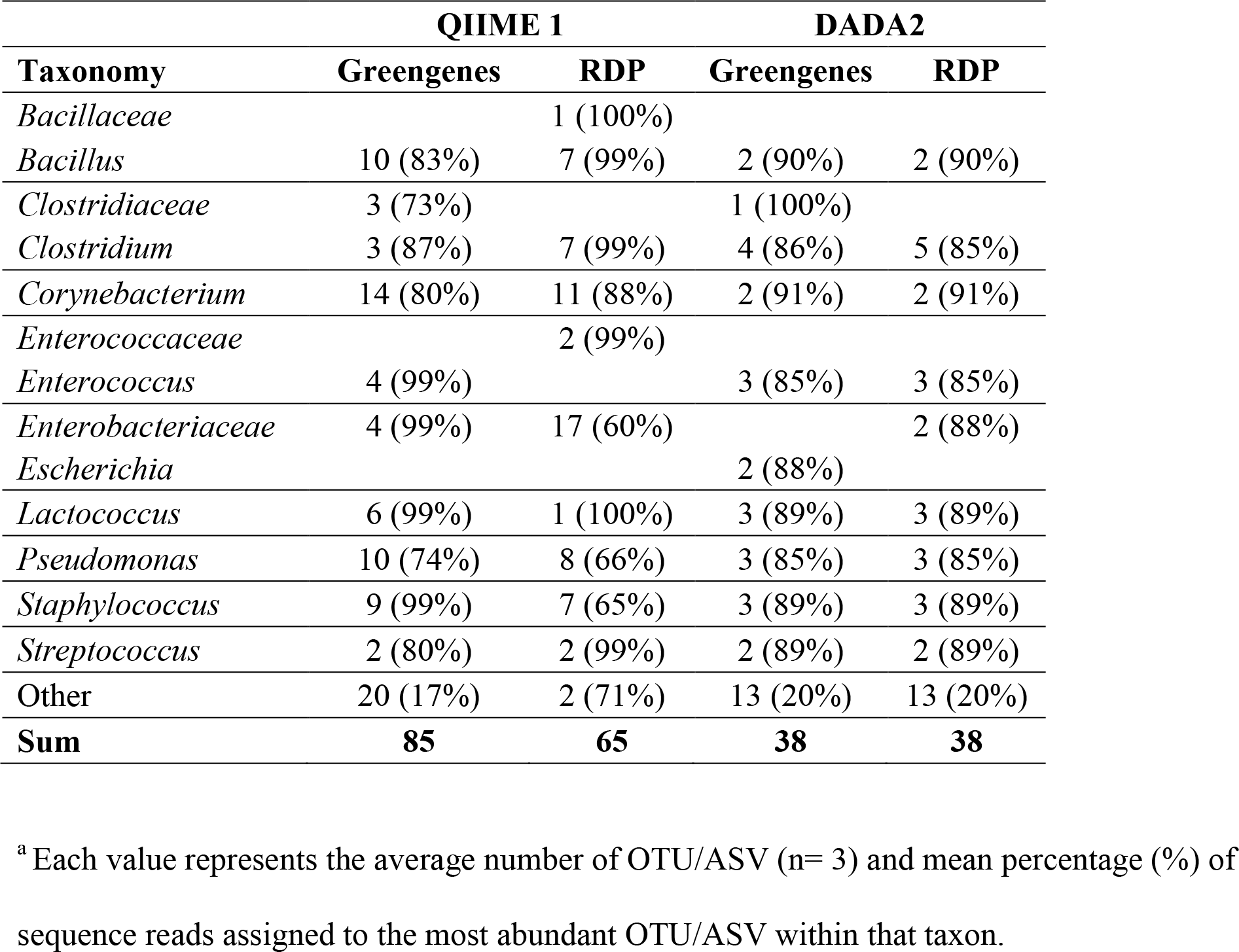
OTU/ASV distribution of the 16S rRNA PCR amplicon mock community following Illumina MiSeq DNA sequencing and paired-end assembly.

| Taxonomy | QIIME 1 |  | DADA2 |  |
| --- | --- | --- | --- | --- |
|  | Greengenes | RDP | Greengenes | RDP |
| *Bacillaceae* |  | 1 (100%) |  |  |
| *Bacillus* | 10 (83%) | 7 (99%) | 2 (90%) | 2 (90%) |
| *Clostridiaceae* | 3 (73%) |  | 1 (100%) |  |
| *Clostridium* | 3 (87%) | 7 (99%) | 4 (86%) | 5 (85%) |
| *Corynebacterium* | 14 (80%) | 11 (88%) | 2 (91%) | 2 (91%) |
| *Enterococcaceae* |  | 2 (99%) |  |  |
| *Enterococcus* | 4 (99%) |  | 3 (85%) | 3 (85%) |
| *Enterobacteriaceae* | 4 (99%) | 17 (60%) |  | 2 (88%) |
| *Escherichia* |  |  | 2 (88%) |  |
| *Lactococcus* | 6 (99%) | 1 (100%) | 3 (89%) | 3 (89%) |
| *Pseudomonas* | 10 (74%) | 8 (66%) | 3 (85%) | 3 (85%) |
| *Staphylococcus* | 9 (99%) | 7 (65%) | 3 (89%) | 3 (89%) |
| *Streptococcus* | 2 (80%) | 2 (99%) | 2 (89%) | 2 (89%) |
| Other | 20 (17%) | 2 (71%) | 13 (20%) | 13 (20%) |
| **Sum** | **85** | **65** | **38** | **38** |

<sup>a</sup> Each value represents the average number of OTU/ASV (n= 3) and mean percentage (%) of sequence reads assigned to the most abundant OTU/ASV within that taxon.

The numbers of ASVs identified with DADA2 from Illumina paired-end assemblies were lower than the numbers of OTUs assigned with QIIME 1 (Table 2). Because DADA2 assigns ASVs independently from taxonomic reference databases, total ASV numbers were the same using both Greengenes and RDP. The only distinction was that one *Clostridiaceae* ASV and both *Escherichia* ASVs identified using Greengenes were designated as *Clostridium* and *Enterobacteriaceae* in RDP (Table 2).

### Illumina MiSeq unassembled, single-end reads

Without read assembly, the QIIME 1 pipeline resulted in 68 and 36 total OTUs with the Greengenes and RDP databases, respectively (Table 3). These OTU numbers were lower compared to the paired-end assemblies (Table 2). DADA2, on the other hand, resulted in a slightly higher number of ASVs (40 ASVs) than the paired-end assemblies (38 ASVs) (Table 2 and Table 3). More reads were regarded as “other” taxa, and this result was most likely due to the shorter lengths of the unassembled, single-end MiSeq reads. Between the two reference databases, Greengenes resulted in more accurate taxonomic assignments with DADA2. Two *Bacillales* ASVs and one *Clostridiales* ASV that were included in the “other” ASV category by RDP were assigned as *Bacillus* and *Clostridiaceae* with Greengenes. Moreover, the *Escherichia/Shigella* ASV in RDP was unambiguously allotted to the *Escherichia* genus by Greengenes (Table 3).

**Table 3.**
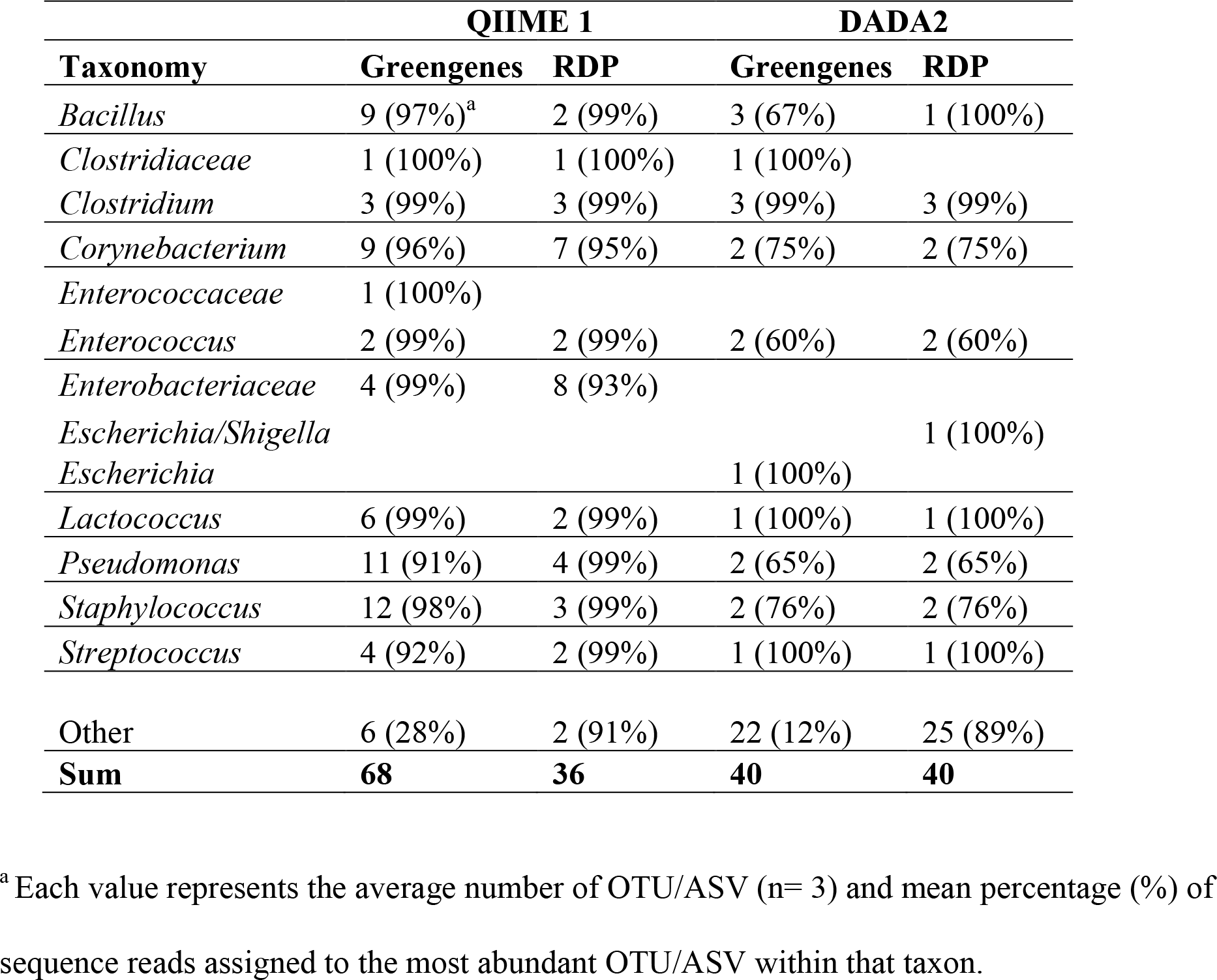
OTU/ASV distribution of the 16S rRNA PCR amplicon mock community following Illumina MiSeq DNA sequencing without paired-end assembly.

### Ion Torrent PGM reads

The application of QIIME 1 with the Greengenes database to the Ion Torrent reads resulted in the highest number of OTUs out of any of the methods applied (Table 4). The use of RDP in QIIME 1 also yielded high OTU numbers, comparable to those found for the paired-end Illumina MiSeq assemblies (Table 4). Conversely, DADA2 resulted in only 13 ASVs (Table 4). The 4 additional ASVs compared to expected were created due to errors in the homopolymer regions (Fig. S1) and the 13 ASVs were distributed across the 9 bacterial taxa included in the mock community, with the exception of one ASV with ambiguous taxonomy (*Bacillales*) identified using RDP, which was identified as *Bacillus* using Greengenes. Greengenes also improved the assignment of *E. coli* to the genus level as opposed to the family level in RDP (Table 4).

**Table 4.** OTU/ASV distribution of the 16S rRNA PCR amplicon mock community following Ion Torrent PGM sequencing.

| Taxonomy | QIIME 1 |  | DADA2 |  |
| --- | --- | --- | --- | --- |
|  | Greengenes | RDP | Greengenes | RDP |
| *Bacillaceae* | 3 (88%)<sup>a</sup> |  |  |  |
| *Bacillus* | 21 (75%) | 8 (95%) | 2 (95%) | 1 (100%) |
| *Clostridium* | 5 (70%) | 7 (99%) | 1 (100%) | 1 (100%) |
| *Corynebacterium* | 15 (75%) | 11 (88%) | 1 (100%) | 1 (100%) |
| *Enterococcaceae* |  | 2 (50%) |  |  |
| *Enterococcus* | 9 (98%) | 2 (99%) | 1 (100%) | 1 (100%) |
| *Enterobacteriaceae* | 13 (95%) | 17 (60%) |  | 1 (100%) |
| *Escherichia* |  |  | 1 (100%) |  |
| *Lactococcus* | 6 (99%) | 1 (100%) | 2 (99%) | 2 (99%) |
| *Pseudomonas* | 13 (71%) | 8 (66%) | 2 (99%) | 2 (99%) |
| *Staphylococcus* | 21 (97%) | 7 (65%) | 2 (99%) | 2 (99%) |
| *Streptococcus* | 6 (83%) | 2 (99%) | 1 (100%) | 1 (100%) |
| Other | 6 (40%) | 2 (71%) | 0 | 1 (100%) |
| **Sum** | **118** | **67** | **13** | **13** |

<sup>a</sup> Each value represents the average number of OTU/ASV (n= 3) and mean percentage (%) of sequence reads assigned to the most abundant OTU/ASV within that taxon.

For each of the three DNA sequencing/read curation methods tested, DADA2 assigned fewer ASVs per taxon and resulted in fewer spurious ASVs than QIIME 1 (UCLUST) assigned OTUs except in Illumina single-end results analyzed with RDP database. DADA2 taxonomic identification was more specific with the Greengenes than the RDP database. Therefore, the combined DADA2/Greengenes approach was used for the subsequent analyses described below.

### Assessments of the gDNA mock community were altered depending on DNA sequencing platform

A gDNA mock community was prepared by mixing equal quantities of gDNA from the nine milk-associated bacterial species prior to barcoded 16S rRNA V4 region PCR amplification. The PCR products were then used for sequencing on either the Illumina MiSeq or Ion Torrent PGM followed by analysis with the DADA2/Greengenes method. More chimeras were found for the gDNA mock community (ranging from 0.3 to 4.6 %) than the PCR amplicon mock community (Table S1). However, except for the known variation in platform-dependent read lengths, nucleotide sequences of the most abundant ASVs assigned to each of the nine mock community species were identical between the Illumina MiSeq (single– and paired-ends) and Ion Torrent PGM platforms (Illumina MiSeq paired-end assembly, Fig. S2; Illumina MiSeq singleend reads, Fig. S3; and Ion Torrent PGM reads, Fig. S4). Nucleotide sequence alignments of those ASVs to the corresponding ASVs identified from the PCR amplicon mock community also showed 100% nucleotide sequence conservation (Illumina MiSeq paired-end assembly, Fig. S2; Illumina MiSeq single-end reads, Fig. S3; and Ion Torrent PGM reads, Fig. S4).

For both Illumina MiSeq assembled and unassembled (single-end) reads, the gDNA mock community resulted in high numbers of ASVs (Table 5). These numbers were higher than found for the PCR amplicon mock community (Table 2 and Table 3). This result was primarily due to the higher quantities of spurious ASVs (e.g. *Anaeroplasma, Bacteroides, Desulfovibrio* and *Prevotella*) (Table S2) present at low proportions. Conversely, the same number of 13 ASVs were found for the gDNA and PCR amplicon mock communities when the Ion Torrent PGM was used (Table 5).

**Table 5.** ASV distribution of the 653 gDNA mock community following different sequencing methods^a^.

| **Taxonomy** | **Illumina paired** | **Illumina single** | **Ion Torrent** |
| --- | --- | --- | --- |
| *Bacillus* | 4 (86%)<sup>b</sup> | 2 (96%) | 2 (95%) |
| *Clostridiaceae* | 3 (92%) | 1 (100%) |  |
| *Clostridium* | 4 (80%) | 2 (94%) | 1 (100%) |
| *Corynebacterium* | 2 (96%) | 1 (100%) | 1 (100%) |
| *Enterococcus* | 3 (88%) | 1 (100%) | 1 (100%) |
| *Escherichia* | 2 (96%) | 1 (100%) | 1 (100%) |
| *Lactococcus* | 3 (88%) | 3 (67%) | 2 (99%) |
| *Pseudomonas* | 3 (94%) | 1 (100%) | 2 (99%) |
| *Staphylococcus* | 2 (90%) | 2 (99%) | 2 (99%) |
| *Streptococcus* | 3 (90%) | 2 (99%) | 1 (100%) |
| Other | 116 (12%) | 111 (10%) | 0 |
| **Sum** | 145 | 127 | 13 |

<sup>a</sup> Results were based on DADA2 analysis pipeline with the Greengenes database.
<sup>b</sup> Each value represents the average number of OTU/ASV (n= 3) and mean percentage (%) of sequence reads assigned to the most abundant OTU/ASV within that taxon.

The alpha diversity indices Shannon Index and Chao1 of the gDNA mock community were elevated compared to the PCR amplicon mock community for both paired-end and singleend Illumina MiSeq results (Fig. 1). Comparisons to *in silico* reconstructions of the mock communities showed that the Shannon Index and Chao1 were also significantly higher than expected for both Illumina methods. For the PCR amplicon mock community, both indices were significantly higher than expected (Fig. 1A and C). However, only Shannon, and not Chao1 reached significance for the gDNA mock community (Fig. 1B and D). On the other hand, when the Ion Torrent PGM was used, the Shannon Index and Chao I of the PCR amplicon and gDNA mock communities resembled expected values (Fig. 1).

**FIG 1.**
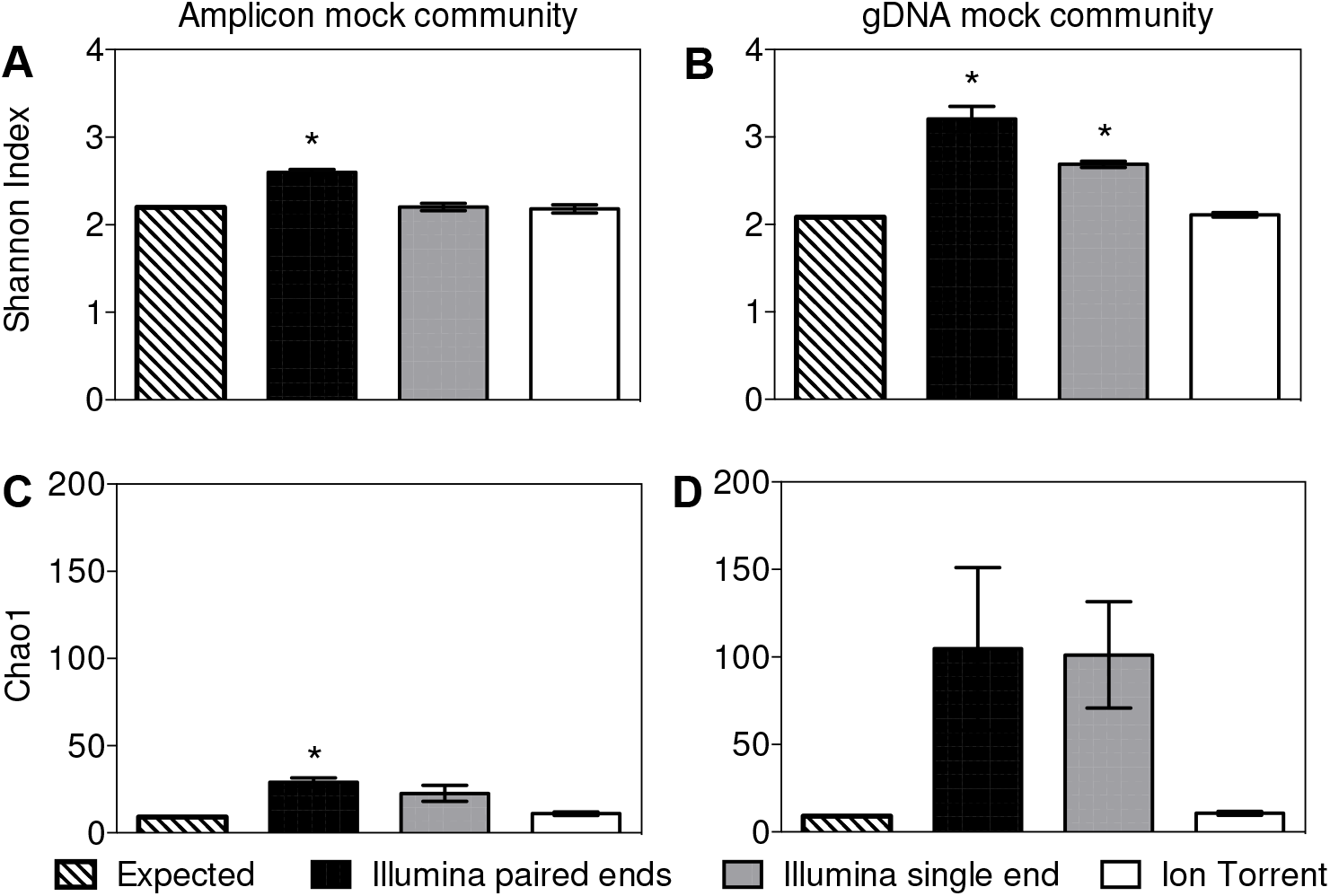
Alpha diversity measurements of mock community samples. Shannon index of (A) PCR amplicon mock community and (B) gDNA mock community. Chaol index of (C) amplicon mock community and (D) gDNA mock community. Results shown were analyzed following DADA2 pipeline and Greengenes database. Each bar represents the mean ± stdev for three replicates. Alpha diversity measurements for each community were compared to expected values using the ANOVA with the Bonferroni’s multiple comparisons test. P values <0.05 were considered to be significant from the expected values and were indicated by the presence of asterisk signs above each bar plot.

### Ion Torrent PGM sequencing with the DADA2/Greengenes method resulted in more accurate representations of the gDNA and PCR amplicon mock communities

DNA sequencing approaches were next compared for their capacity to yield the expected beta diversity and proportions of bacterial taxa included in the two mock communities. According to UPGMA hierarchical clustering of Bray-Curtis dissimilarity metrics, results from the three DNA sequencing approaches (Illumina MiSeq paired-end assembly, single-end, and Ion Torrent PGM) formed separate clusters away from the expected bacterial composition, independent of the gDNA or 16S rRNA PCR amplicon community type (Fig. 2A). Conversely, UPGMA of the weighted Unifrac distance metrics, clustered the sequences according to mock community type (Fig. 2B). These comparisons showed that the gDNA mock communities sequenced with the Ion Torrent PGM were the most similar to theoretical (expected) proportions. No single method was found best suited for representing the PCR amplicon mock community (Fig. 2B). To assess whether the use of DADA2/Greengenes influenced this outcome, the other data analysis methods were compared and it was found that the DNA sequencing platform used was consistently influential on mock community beta-diversity (e.g. QIIME 1 with the Greengenes database is shown in Fig. S5).

**FIG 2.**
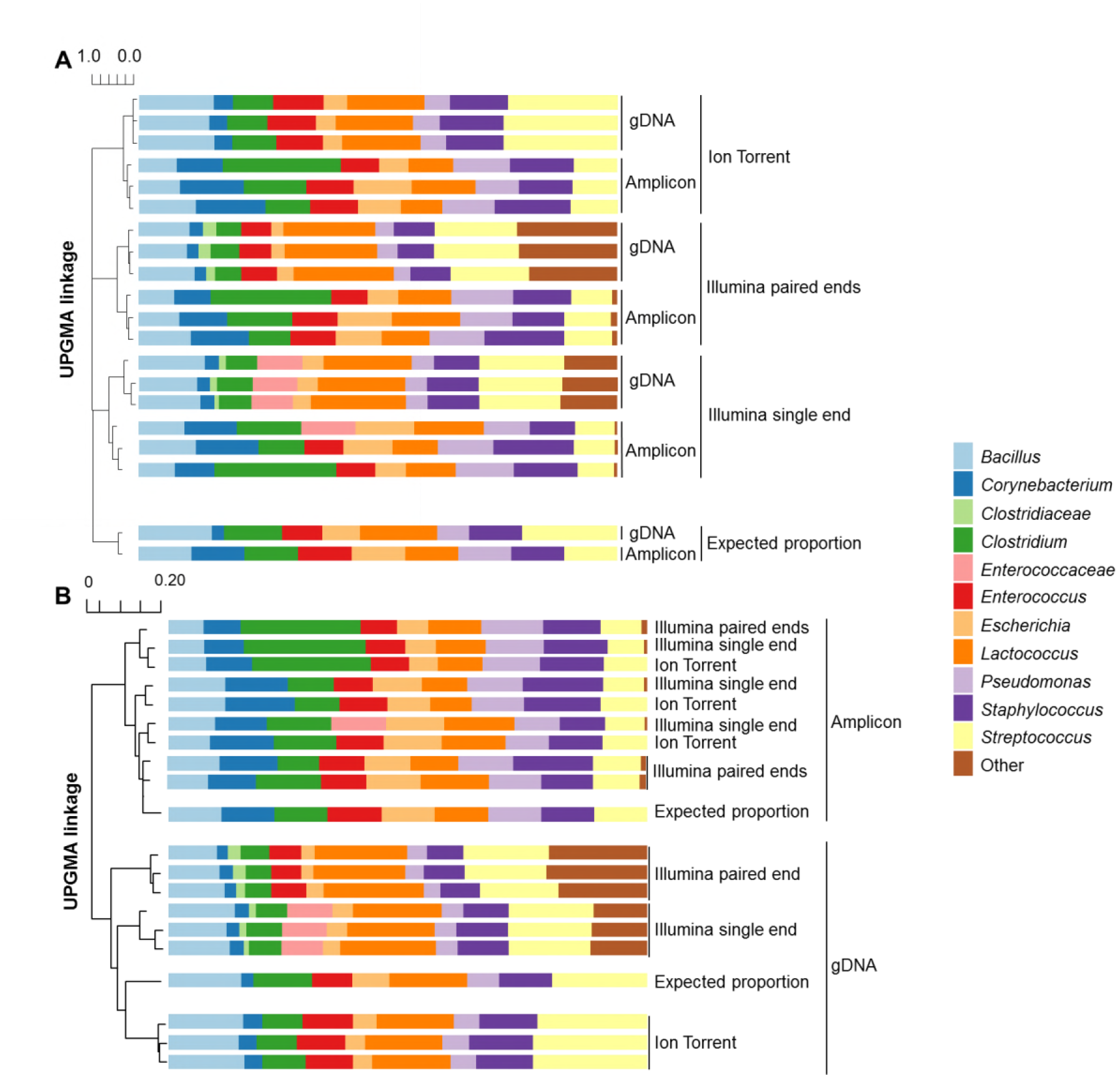
Relative proportions of taxa and UPGMA hierarchical clustering of the mock communities. UPGMA hierarchical clustering was based on the (**A**) Bray Curtis dissimilarity matrix and (**B**) weighted Unifrac distance matrix. Expected taxa (9 bacterial species) are labelled with the corresponding taxonomic level from the DNA sequencing results. Each bar contains the results from each of the three mock community replicates tested using different DNA sequencing methods. Results shown were analyzed following the DADA2 pipeline with Greengenes database.

We next compared the relative abundances of individual taxa across the three DNA sequencing approaches. For the 16S rRNA PCR amplicon mock community, the proportions of most bacterial species were mostly not significantly altered compared to expected theoretical values. Exceptions to this finding were the reduced proportions of *Enterococcaceae* and *Enterococcus* found for Illumina paired-end assemblies and *Streptococcus* for both Illumina MiSeq methods as well as the Ion Torrent PGM platform (Fig. 3). For the gDNA mock community, the proportions of *Escherichia* and *Streptococcus* were significantly different from expected for all three DNA sequencing platforms. The proportions of *Pseudomonas* were also significantly lower than expected for the single- and paired-end assemblies from the Illumina MiSeq, and the proportions of *Bacillus* and *Lactococcus* were also altered for the paired-end assemblies. Lastly, there were higher proportions of “other” taxa for both Illumina MiSeq methods (Fig. 3). Overall, even though DNA sequencing with the Ion Torrent PGM combined with DADA2/Greengenes analysis did not completely provide the expected bacterial composition, this approach resulted in the most accurate representations of the bacteria and their proportions in both gDNA and PCR amplicon mock communities.

**FIG 3.**
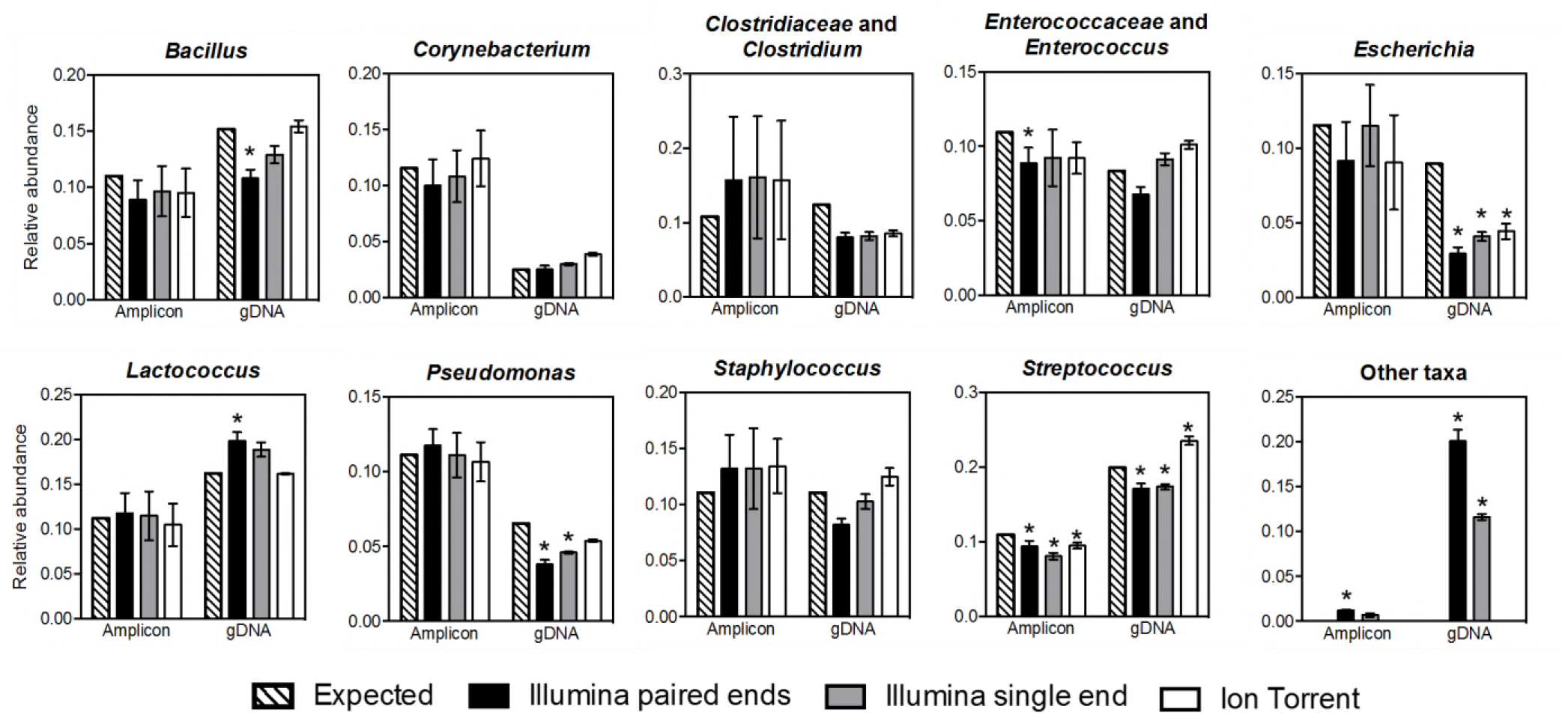
Relative abundance of taxa in the 16S rRNA PCR amplicon and gDNA mock communities. Relative abundance of expected taxa are labelled with the corresponding taxonomic level from sequencing results. Amplicon stands for the 16S rRNA PCR amplicon mock community and gDNA stands for the gDNA mock community. Results shown were analyzed following DADA2 pipeline with Greengenes database. Each bar represents the mean ± stdev for three replicates. Proportions for each community were compared to expected proportions using the ANOVA with the Bonferroni’s multiple comparisons test. P values <0.05 were considered to be significant from the expected values and were indicated by the presence of asterisk signs above each bar plot.

## Discussion

By comparing DNA sequencing methods, analysis algorithms, and reference databases using dairy relevant bacterial DNA (PCR amplicon and gDNA) mock communities, we found that the DADA2/Greengenes data analysis methods with the Ion Torrent PGM yielded the most accurate interpretations of the 16S rRNA V4 variable region of dairy mock communities relative to the other methods (Illumina MiSeq, QIIME 1, RDP) tested.

Application of DADA2 and QIIME 1 (UCLUST) analysis pipelines to the same 16S rRNA gene data showed that DADA2 assigned fewer total and spurious OTUs/ASVs than QIIME 1 even with stringent filtering (32). Because read length was kept consistent within each DNA sequencing and data assembly platforms, platform-specific differences in OTU/ASV numbers were mainly derived from the core algorithms used for filtering and clustering representative sequences. The DADA2 core algorithm includes the error-rate-based denoising, the isBimeraDenovo chimera identification, and the ASV inference (26). In QIIME 1, the core analysis includes the USEARCH chimera identification and OTU picking strategy (33). Comparison of OTUs and ASVs using the DADA2 and QIIME 1 pipelines, respectively, also showed that the DADA2 analysis pipeline was able to assign ASVs to more specific taxonomic levels (genus) than QIIME 1. This could be the result of the different taxonomy classifiers employed by DADA2 (RDP’s naive Bayesian classifier) and QIIME 1 (UCLUST classifier) (34).

OTU and ASV taxonomy assignments were also compared with consideration to 16S rRNA gene reference databases. Results from the different combinations of analysis methods and reference databases showed that the majority of OTUs/ASVs detected were representatives of bacterial taxa included in the 16S rRNA PCR amplicon and gDNA mock communities. Each bacterial species was represented by at least a single OTU/ASV. Additional OTUs/ASVs were largely due to low abundance sequence variants. DNA sequences of the predominant ASVs/OTUs were 100% identical between gDNA and PCR amplicon mock communities, further supporting the precision of the technique. When QIIME 1 was applied, the RDP gold database (35) yielded lower numbers of total OTUs than found with Greengenes 13.8, independent of whether the Illumina MiSeq or Ion Torrent PGM were used to generate the DNA sequence reads. However, the RDP gold database has not been updated since 2011 (35) and could potentially be missing many bacterial sequences, leading less differentiation between OTUs. With the DADA2 pipeline, ASVs were inferred prior to taxonomy assignment (26), resulting in the same total ASV numbers for both RDP 11.5 and Greengenes 13.8 databases. However, assignments of DADA2 ASVs were still influenced by reference database-specific taxonomic nomenclature and DNA sequences (36) such that Greengenes was provided deeper, more accurate taxonomic assignments than found with RDP.

The Illumina MiSeq and Ion Torrent PGM methods also clearly impacted the outcomes of our mock community analyses. The Illumina MiSeq is well-established and known for its low error rate, high volume read outputs, and low sequencing cost per Gb (37, 38). The Ion Torrent PGM method is recognized for shorter run times, lower instrument cost, and flexibility in sequencing scale per run by the use of different sequencing chips (37, 38), and therefore could be of relevance for dairy and other food products with short shelf-life times and low sample numbers. For both single-end and paired-end assembled Illumina MiSeq reads, a greater number of unexpected taxa and OTUs/ASVs were observed compared to the Ion Torrent PGM. This finding could be the result of substitution errors Illumina MiSeq read accumulated leading to taxonomy misidentification (39, 40). This issue became particularly apparent with the finding of unrelated taxa such as *Anaeroplasma, Bacteroides, Desulfovibrio* and *Prevotella* that were not included in the gDNA mock community. Use of the Ion Torrent PGM with our read trimming parameters resulted in the lowest numbers of DADA2 assigned ASVs. At 13 ASVs for both mock communities, this number was only slightly greater than the nine predicted. All thirteen ASVs were repeatedly assigned to members of the mock communities, except for one low abundance ASV when RDP was applied. The four additional ASVs were the result of read errors in the homopolymer regions, a common Ion Torrent error model (20, 37, 38). Interestingly, the Ion Torrent PGM reads resulted in the highest numbers of OTUs when QIIME 1 was used to analyze the data. This might have been due to the higher number of erroneous reads that were passed by QIIME 1 filtering, but were identified as sequence chimeras and artifacts by DADA2.

Alpha and beta diversity metrics were impacted by the DNA sequencing method. Calculations of beta diversity from the DADA2/Greengenes pipeline showed that the results from the Illumina MiSeq were similar to Ion Torrent PGM, especially for the PCR amplicon mock community. However, both paired and single end Illumina MiSeq methods resulted in erroneous estimates of *Streptococcus* and *Lactococcus*. Because *Streptococcus* and *Lactococcus* share similar 16S rRNA gene sequences, variation in their relative abundances could be caused by the accumulation of substitution errors, a common error that occurs with those instruments (39, 40). Alpha-diversity and beta-diversity assessements on data collected with the Ion Torrent PGM and analyzed by DAD2/Greengenes most closely resembled the *in silico* reconstructions of the mock communities. Taxonomic proportions were also less impacted compared to results from the Illumina MiSeq and limited to reduced proportions of *Escherichia* and increased *Streptococcus* levels. This finding is similar to previous studies in which *Proteobacteria* proportions were underestimated and *Firmicutes* proportions were overestimated due to negative correlations with genomic G+C content (41). To minimize PCR preference of A+T rich bacterial organisms, methods such as increasing the denaturing time (41) could be used for future dairy microbiome studies.

By the use of bacterial DNA standards from nine dairy-relevant bacterial species, we found that DNA sequencing and analysis pipelines contributed significant variations to OTU/ASV distributions and observed bacterial diversities. Moreover, PCR biases and errors from multi-template DNA amplifications are not entirely filtered with the Illumina MiSeq method. Overall, the Ion Torrent PGM DNA sequencer combined with the DADA2/Greengenes pipeline led to more accurate OTU/ASV assignments and bacterial diversity measurements under our study conditions. With DADA2 being wrapped in the QIIME 2, we agree with the QIIME 2 developers that new sequencing results should be analyzed using QIIME 2 with standardized analysis pipeline (e.g: DADA2) instead of QIIME 1 (UCLUST) (42). Further improvements might be reached by refinements to taxonomy classifiers (34) and updating reference databases to emphasize bacteria found in different environments such as dairy foods (36). Moreover, additional studies are required to further establish the relevance of PCR protocols on the bacterial diversity outcomes, and the analysis methods should be tested on mock communities consisting of intact bacterial cells exposed to selected DNA extraction and purification methods (43). The findings here and the continued development of bacterial diversity analysis methods should result in reliable comparisons within and between microbial habitats.

## Materials and Methods

### Bacterial strains and culture conditions

Bacterial strains representing species commonly found in bovine milk were used to construct a mock bacterial community (Table 1). Each bacterial strain was grown in standard laboratory culture medium with negative controls for that species and harvested at early stationary phase by centrifugation at 13,000xg for 2 min. Laboratory culture medium was as follows: *Bacillus subtilis, Pseudomonas fluorescens*, and *Escherichia coli* – LB Lennox broth (Thermo Fisher Scientific); *Enterococcus faecalis* and *Streptococcus agalactiae* – Brain Heart Infusion broth (Thermo Fisher Scientific); *Staphylococcus aureus* – Tryptic Soy broth (Becton Dickinson); *Corynebacterium bovis* – Tryptic Soy broth (Becton Dickinson) with 0.1% tween 80; *Lactococcus lactis* – M17 broth (Becton Dickinson) with 0.5% glucose; and *Clostridium tyrobutyricum* – Reinforced Clostridial broth (Becton Dickinson). All strains were incubated at 37 °C, with the exception of *B. subtilis, L. lactis* and *P. fluorescens*, which were incubated at 30 °C. B. subtilis, *C. bovis*, E. faecalis, E. coli and *P. fluorescens* were grown under aeration (250 rpm).

### Genomic DNA extraction and PCR amplification

Genomic DNA was extracted using the MagMAX Total Nucleic Acid Isolation Kit (Thermo Fisher Scientific, Vilnius, LT) according to the manufacturer’s protocol with the repeat bead beating method on a FastPrep-24 instrument (MP Biomedicals LLC). DNA concentration was measured with the Qubit 3.0 Fluorometer using the Qubit dsDNA HS Assay Kit (Life Technologies, Eugene, OR). PCR amplification was performed using ExTaq DNA polymerase (TaKaRa, Otsu, Japan) and primers F515 and R806 (44) with a random 8 bp barcode on the 5’ end of F515 for sample multiplexing (45, 46). PCR was initiated at 94 °C for 3 min and followed by 35 cycles of 94 °C for 45 sec, 54 °C for 60 sec, and 72 °C for 30 sec with a final extension step at 72 °C for 10 min. Negative controls were run for each barcoded primer. No PCR product for the negative controls was observed on a 1.5% agarose gel. PCR products were pooled and then gel purified with the Wizard SV Gel and PCR Clean-Up System (Promega, Madison, WI, USA).

### Preparation of the mock communities

For the gDNA mock community, 100 ng gDNA isolated from each of the strains was pooled in three separate replicates. The proportion of each bacterial strain in the gDNA mock community was determined by taking into account the genome size and 16S rRNA gene copy number (Table 1). To construct the amplicon mock community, gDNA of the nine bacterial strains was amplified in triplicate by using three different barcoded PCR primers. Amplicon concentrations were measured with the Quant-iT PicoGreen dsDNA Assay Kit (Life Technologies, Eugene, OR) prior to pooling at equal molar concentrations.

### DNA sequencing

For Illumina sequencing, the KAPA HTP library preparation kit (KK8234, Kapa Biosystems, Pittsburgh, PA) was used for the ligation of NEXTflex™ adapters (Bioo Scientific, Austin, TX) to the 16S rRNA amplicons prior to 250 bp paired-end sequencing (with 7% PhiX control) on an Illumina MiSeq instrument at the University of California, Davis genome center (http://genomecenter.ucdavis.edu/). For Ion Torrent sequencing, non-barcoded Ion A and Ion P1 adapter were ligated to the pooled amplicons followed by templating, enrichment and sequencing on the One-Touch 2, One-Touch ES systems and Ion PGM using the 400 sequencing kit and a 318 v2 chip (Life Technologies, Carlsbad, CA).

### 16S rRNA gene sequence analysis

An *in silico* mock community, termed “expected”, was created using the 16S V4 amplicon sequences from published genomes and reference genomes for the specific bacterial species (Table 1). In addition, the expected 16S V4 region copy numbers were normalized based on the genome size and 16S rRNA gene copy numbers.

Illumina MiSeq sequencing outputs were trimmed with the fastx_tools (47) to keep the first 245 and 170 bases for the forward and reverse reads respectively. Ion Torrent sequence output BAM file was converted to FASTQ format using BEDTools (48) and reads shorter than 200bp were also removed. The first 280 bases of the Ion Torrent reads were kept.

The FASTQ files were then analyzed with QIIME version 1.9.1 and DADA2 1.6.0 (26, 49). In QIIME 1, Illumina reads from the two orientations (forward and reverse) were analyzed either with or without assembly where the *join_paired_ends.py (fastq-join* method) (50) script was used with minimum 100 bp overlap and 1% maximum difference between overlapping sequences. Ion Torrent, single-end and paired-end assembled Illumina FASTQ files were then removed of barcode (8 bases) and primer regions (forward primer: 21 bases, reverse primer: 20 bases) and were demultiplexed using the *split_libraries_fastq.py* script with no barcode error and quality filtered at Q30. Chimeric sequences were identified and removed using USEARCH method (33, 35) and the Greengene database version 13.8 (51, 52). Sequences from both Illumina and Ion Torrent as well as the *in silico* mock community with expected proportions were merged together as one fasta file for operational taxonomic unit (OTU) clustering using the *pick_open_reference_otus.py* script with recommended parameters (32) and the UCLUST method at 97% similarity thresholds. Greengenes version 13.8 (51, 52) and RDP_GOLD (35) databases were used as references for OTU assignments. Archaea, Chloroplast and low abundance (0.005%) OTUs were removed from the OTU tables (32).

In DADA2, for single-end analysis, the truncated Illumina and Ion Torrent FASTQ files after barcode (8 bases) and primer sequences (forward primer: 21 bases, reverse primer: 20 bases) trimming were demultiplexed using *split_libraries_fastq.py* script with no barcode error and no quality filter (-r 999, -n 999, -q 0, -p 0.0001). Since the single-end reads were already quality trimmed, no additional truncation was performed in DADA2 to be consistent in read length with QIIME 1 analysis. For paired-end analysis, in order to get matched sequence files, raw Illumina reads were demultiplexed in pairs using the idemp tool (53) with no barcode error. Barcode, forward and reverse primer regions were then trimmed with fastx_tools (47). Resulting reads were truncated in DADA2 to keep the first 196 bases of the forward reads and 121 bases of the reverse reads, which were then merged with 51bp minimum overlap to be consistent with QIIME 1 method in resulting read length. The error model learning, dereplication and chimera removal steps were performed in DADA2 with default parameters. Taxonomy was assigned to the resulting amplicon sequence variants (ASVs) using RDP database version 11.5 (54) and Greengenes database version 13.8 with the minimum bootstrap confidence at 80 (51, 52). Ion Torrent, Illumina single-end and paired-end assembled reads were merged together with the *in silico* mock community using the phyloseq package in R (55), and singletons and low abundance (0.005%) ASVs were removed to be consistent with QIIME 1 analysis.

### Accession numbers

Joined– and single-end DNA sequences after quality filtering and trimming were deposited in the Qiita database (https://qiita.ucsd.edu) under study ID 11351 and in the European Nucleotide Archive (ENA) under accession number ERP104377.

### Statistics

OTU/ASV counts were rarefied at 5483 sequences per sample to retain all samples for downstream analyses. Significant differences in the observed mock community composition (alpha diversity and taxonomic distribution) were determined by ANOVA with the Bonferroni’s multiple comparisons test. A p-value of <0.05 indicates significance. The significance of sample clustering was indicated by permutational multivariate analysis of variance using the adonis function from the vegan package in R (56) with a p-value of <0.05 through 9,999 permutations.

## Acknowledgements

The authors would like to thank Yanin Srisengfa for her extensive help with bacterial culture. The authors would also like to thank NIZO food research for providing strain *Lactococcus lactis* IL1403, and Dr. Steven Lindow at University of California, Berkeley for providing strain *Pseudomonas fluorescens* A506.

The California Dairy Research Foundation grant “Rapid methods for microbial detection and quality assessment of milk” funded this study. The funding agency did not participate in study design, data collection, or interpretation of the data.

